# Validating amino acid variants in proteogenomics using sequence coverage by multiple reads

**DOI:** 10.1101/2022.01.08.475497

**Authors:** L.I. Levitsky, K.G. Kuznetsova, A.A. Kliuchnikova, I.Y. Ilina, A.O. Goncharov, A.A. Lobas, M.V. Ivanov, V.N. Lazarev, R.H. Ziganshin, M.V. Gorshkov, S.A. Moshkovskii

## Abstract

Mass spectrometry-based proteome analysis usually implies matching mass spectra of proteolytic peptides to amino acid sequences predicted from nucleic acid sequences. At the same time, due to the stochastic nature of the method when it comes to proteome-wide analysis, in which only a fraction of peptides are selected for sequencing, the completeness of protein sequence identification is undermined. Likewise, the reliability of peptide variant identification in proteogenomic studies is suffering. We propose a way to interpret shotgun proteomics results, specifically in data-dependent acquisition mode, as protein sequence coverage by multiple reads, just as it is done in the field of nucleic acid sequencing for the calling of single nucleotide variants. Multiple reads for each position in a sequence could be provided by overlapping distinct peptides, thus, confirming the presence of certain amino acid residues in the overlapping stretch with much lower false discovery rate than conventional 1%. The source of overlapping distinct peptides are, first, miscleaved tryptic peptides in combination with their properly cleaved counterparts, and, second, peptides generated by several proteases with different specificities after the same specimen is subject to parallel digestion and analyzed separately. We illustrate this approach using publicly available multiprotease proteomic datasets and our own data generated for HEK-293 cell line digests obtained using trypsin, LysC and GluC proteases. From 5000 to 8000 protein groups are identified for each digest corresponding to up to 30% of the whole proteome coverage. Most of this coverage was provided by a single read, while up to 7% of the observed protein sequences were covered two-fold and more. The proteogenomic analysis of HEK-293 cell line revealed 36 peptide variants associated with SNP, seven of which were supported by multiple reads. The efficiency of the multiple reads approach depends strongly on the depth of proteome analysis, the digesting features such as the level of miscleavages, and will increase with the number of different proteases used in parallel proteome digestion.

**Graphical abstract:** 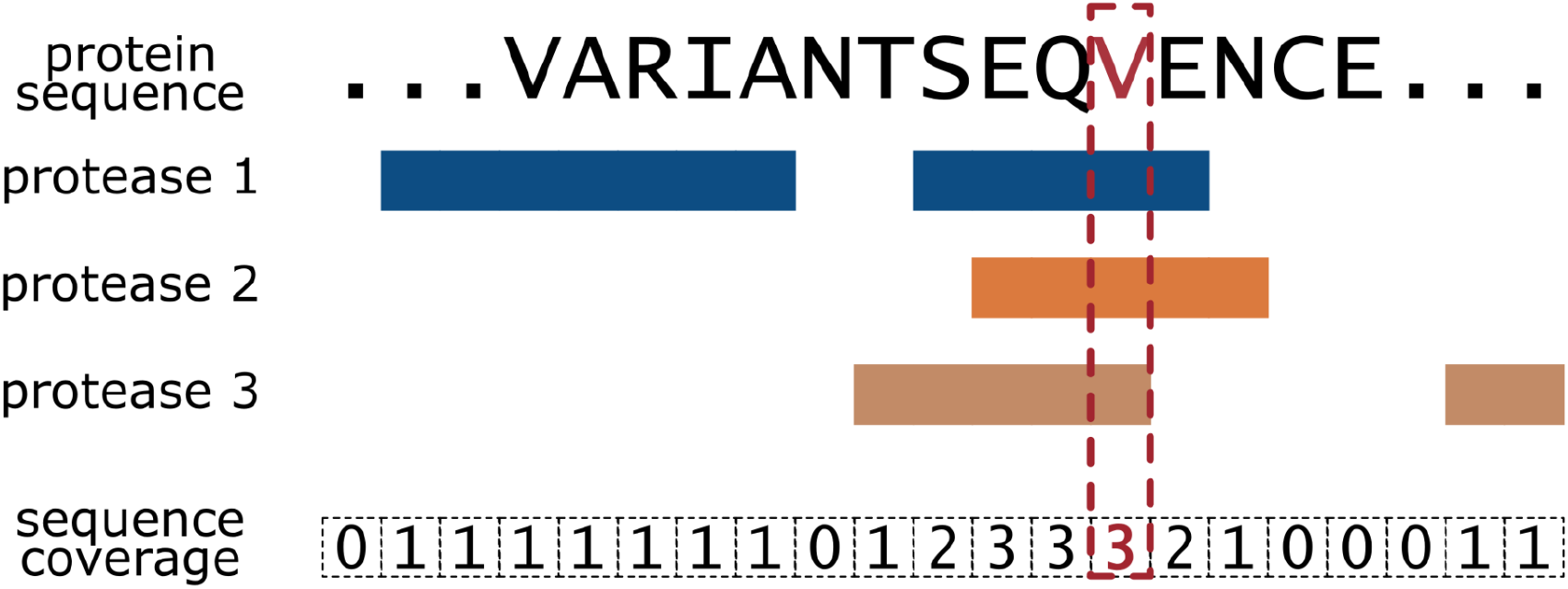

## 1. Introduction

A majority of mass spectrometric studies of proteomes currently use the bottom-up, also called shotgun, technique [1]. The key part of the bottom-up workflow is protein digestion by a protease to produce a peptide mixture which is subject to the follow-up liquid chromatography/tandem mass spectrometry (LC-MS/MS) analysis. Peptides are much simpler macromolecules for identification from both LC and MS/MS viewpoints compared with proteins, yet, this approach complicates subsequent protein identification based on the peptide-spectrum matches (PSM), posing the so-called protein inference problem [2,3]. The most effective way to attribute a measured tandem mass spectrum is by means of a genomic database, predicting a limited set of peptides from the theoretical proteome. An algorithm will select the best match to any eligible mass spectrum. To filter out unreliable predictions, the false discovery rate (FDR) is then calculated based typically on the so-called target-decoy approach (TDA) [4]. The PSMs passing the FDR threshold of, typically, 1%, constitute the list of identified peptides used further for protein inference.

Although this practice has been challenged by bioinformaticians [5], protein identifications are generally considered much more reliable if they are supported by multiple distinct peptide identifications. Otherwise, if one reports a new finding based on a single peptide identified by one or more PSMs, some additional validation is usually required. Unfortunately, this is a typical story in proteogenomics, in which important single peptide-based reports include variants predicted from genetic polymorphisms [6], alternative splicing events [7], RNA editing [8], or so-called missing proteins [9].

The above cases are typically resolved by targeted proteomic methods, such as multiple/selected reaction monitoring (MRM/SRM) or parallel reaction monitoring (PRM) [10]. For most applications, where amino acid variants to be detected originate from actionable or neoantigenic mutations, one can use targeted methods directly, omitting the shotgun proteome analysis step; however, this approach requires a large cohort of synthetic peptide standards and a lot of instrumentation time, further limiting its utility for wider clinical applications.

Several orthogonal methods have been introduced to increase reliability of peptide identifications by random sampling in shotgun proteomics. Addition of specific peptide features is used for scoring PSMs, such as a chromatographic retention time [11], intensity pattern in fragmentation spectra [12–14], etc. Rapidly accumulating data from large-scale studies made it possible to explore deep learning approaches for efficient prediction of these features to further enhance identification of PSMs. At the same time, there are still no standards in the field for validation of single peptide-based findings and further improvements to this end may be suggested.

For applications of shotgun proteomics such as proteogenomics, where single amino acid polymorphisms and, correspondingly, single peptide identification is often the only available option, it is desirable to improve reliability of these identifications within the scope of the method, ideally, avoiding additional procedures, such as targeted mass-spectrometry. Elaborating further on the similarities between shotgun proteomics and next generation sequencing (NGS) of nucleic acids, which is also a shotgun approach, note that single nucleotide polymorphisms are confirmed by multiple overlapping reads in NGS [15]. The idea of this work is to use the same strategy for shotgun proteomics data. Overlapping reads in NGS are intrinsic for many methods of sequencing [16]. In a similar fashion, one can define the ‘read’ in shotgun proteomics. It is, naturally, a single PSM, which is a hypothesis that a given mass spectrum is produced by a given peptide compound listed in a genomic database, with some measure of reliability [17]. PSMs from different mass spectra and with different retention times and other measurable parameters are formally independent. Thus, if two or more different PSMs map to overlapping parts of a protein sequence, the corresponding identification of the overlap becomes increasingly more reliable. A common source of overlapping PSMs in shotgun proteomics is identification of a properly cleaved tryptic peptide and its miscleaved counterpart. If, for example, a sequence variant of interest would fall into the overlapping part, we must think that it is identified more reliably than from a single PSM. A hunt for overlapping reads in shotgun proteomics is the rationale behind the approach explored in this work.

The embodiment of this idea is focused primarily on the data dependent acquisition (DDA) mode as the concept of peptide/PSM FDR for data independent acquisition (DIA) has yet to be established and such notations as the group-specific FDR [18] are not available. At the same time, for many DIA applications, DDA runs are also made to attribute identities to peptides by the conventional target decoy-method and then DIA provides better quantification of proteins being not focused on single amino acid variants [6].

Controlling the efficiency of trypsinolysis to intentionally produce overlapping cleaved and miscleaved peptides does not seem feasible due to different kinetics of the cleavage process for different protein sequences [19]. An easier way to obtain overlapping reads is the use of multiple proteases known in the field [20–23]. In this work, we analyzed publicly available datasets and generated our own experimental data for the model HEK-293 cell line using multiple proteases to conceptualize the approach of utilizing the sequence coverage by multiple reads for validation of single amino acid variants.

## 2. Experimental procedures

### 2.1. Cell culture

Cell culture was managed in accordance with our previous work [24]. Briefly, HEK-293 cells were obtained from ATCC (accession number CRL-1573; Manassas, VA). The 40th passage of HEK-293 was used for proteome analysis.

Cryogenically preserved cells were thawed and expanded in culture medium (DMEM) supplemented with 10% (w/v) fetal bovine serum (FBS) and 100 units/mL gentamicin (all from Gibco, Thermo Fisher Scientific, Bremen, Germany) in a humidified CO2 incubator under standard conditions (5% CO_2_, 37 °C). The medium was exchanged every 2 days. To prepare cell samples for protein extraction, the cells were detached with 0.05% Trypsin-EDTA solution (PanEko, Moscow, Russia), washed 3 times with PBS, and counted. Aliquots of the resulting cell suspension were centrifuged, the supernatant was removed, and cells were frozen in liquid nitrogen. The cell pellets were kept frozen in liquid nitrogen vapor until use.

### 2.2. Cell lysis and protein digestion

Cell pellets of one million cells each were resuspended in lysis buffer containing 4% SDS in 100 mM TEABC, pH 8.5, incubated for 5 minutes and subjected to sonication by Qsonica Q55 ultrasonic homogenizer (Qsonica, USA) at 70% amplitude using 10 series of 10 one-second-duration impulses. After that, the samples were incubated for 10 minutes at 85 °C and centrifuge for 10 minutes at 16000 × g. Later on, protein disulfide bonds were reduced by addition of dithiothreitol (DTT) up to 5 mM and incubation for 30 minutes at 56 °C and the cysteine residues were alkylated with chloroacetamide (CAM) in the final concentration of 10 mM for 15 minutes at room temperature at dark.

At the next step, the proteins were precipitated with cold acetone. First, we precipitated the proteins from 10 μL of the lysate to measure protein concentration after the precipitation. Next, the volume containing 30 μg of total protein was taken from each sample for the future analysis.

For precipitation, 4 volumes of cold (−20 °C) acetone were added to the samples. The samples were vortexed and left for 120 minutes at −20 °C to let the proteins thoroughly precipitate. After the incubation period, the samples were centrifuged for 10 minutes at 13,000-15,000 × g at 4 °C followed by rinsing the pellet with cold acetone twice without mixing. Then the acetone was carefully removed and the tubes were left with the lids open for 30 minutes to let the pellets dry up.

For digestion, 30 μL of each protease, namely trypsin (Promega Gold), LysC and GluC (all from Promega, USA) at a concentration of 0.02 μg/μl in 50 mM TEABC were added straight to the pellets. The final w/w proportion of each protease to total protein was 1:50. Then the samples were incubated overnight at 37 °C. To stop the reaction, trifluoroacetic acid (TFA) up to 1% (v/v) was added to each tube.

### 2.3. Peptide desalting and clean-up

For peptide desalting and clean-up, in-house made stage tips containing SDB-RPS membrane (Empore-3M, CDS Analytical, USA) were used. The tips were prepared as described earlier [25] with the use of 3 pieces of membrane in each tip. The samples were loaded into the tips and the tips were centrifuged at 1200 rpm (about 70 × g) in the BioSan Multi-spin MSV-6000 centrifuge until the solution had passed through the membrane. At the next step, washing with 100 μL 0.2% TFA was performed at the same speed. The peptides were eluted by passing 60 μL of 70% acetonitrile (ACN) with 5% ammonia through the tips into the clean tubes at the speed as low as 1000 rpm (about 50 × g) in the same centrifuge. The peptide samples were dried up in the vacuum concentrator (Labconco, USA).

### 2.4. Liquid chromatography and mass spectrometry

For the LC-MS analysis, the samples were reconstituted in 0.1% TFA and loaded onto an Acclaim PepMap 100 C18 (100 mm × 2 cm) trap column (Thermo Fisher Scientific, USA) in the loading mobile phase (2% ACN, 98% H_2_O, 0.1% TFA) at 10 mL/min flow and separated at 40 °C on a 75 mm × 50 cm Acclaim PepMap 100 C18 LC column (Thermo Fisher Scientific, USA) with particle size 2 mm. Reverse-phase chromatography was performed with an Ultimate 3000 Nano LC System (Thermo Fisher Scientific), which was coupled to the Orbitrap QExactive HF mass spectrometer via a nano electrospray source (Thermo Fisher Scientific). Water containing 0.1% (v/v) formic acid (FA) was used as a mobile phase A and ACN containing 0.1% FA (v/v), 20% (v/v) H_2_O as a mobile phase B. Peptides were eluted from the trap column with a linear gradient: 3–35% solution B for 105 min; 35–55% B for 18 min, 55–99% B for 0.1 min, 99% B during 10 min, 99–2% B for 0.1 min at a flow rate of 300 nL/min. After each gradient, the column was reequilibrated with the phase A for 10 min. MS data was collected in DDA mode (TopN = 15). MS1 parameters were as follows: 120K resolution, 350–1400 scan range, max injection time 50 msec, AGC target 3×10^6^. Ions were isolated with 1.2 m/z window, preferred peptide match and isotope exclusion. Dynamic exclusion was set to 30 s. MS2 fragmentation was carried out at 15K resolution with HCD collision energy 28, max injection time – 80 msec, AGC target – 1×10^5^. Other settings were as follows: charge exclusion - unassigned, 1, 6–8, >8.

### 2.5. MS/MS data processing and coverage calculation

All raw data were converted to mzML format using Proteowizard’s MSConvert [26] and searched using IdentiPy v0.3.4 [27] with search parameters specified below. Search results were post-processed with Scavager v0.2.9 [28], used both to increase sensitivity at 1% FDR and to perform the merging of search results across replicates and fractions as needed. Additional reanalysis was performed by merging results across different proteases to produce summary statistics, such as the total number of peptides and proteins identified at 1% FDR, also using Scavager. Proteome coverage calculations were performed using in-house Python scripts.

Human melanoma cell line 82 data [29] were searched against the Swissprot human database. Precursor and fragment ion mass tolerances were optimized using the auto-tune feature of IdentiPy; carbamidomethylation of cysteine was set as a fixed modification, and oxidation of methionine was allowed as a variable modification. Tryptic digestion rule was specified, with full specificity and up to two miscleavages allowed. Other data sets mentioned in Table 1 were processed using the same settings, except that cleavage on arginine and lysine was specified for samples where LysC was added to trypsin for digestion. Human brain data [30] were processed as above, except fragment ion mass tolerance was set at 0.01 Da and precursor ion tolerance was set at 10 ppm (no auto-tune). For LysC digestion, the cleavage rule was set to LysC and up to two miscleavages were allowed. Search parameters for Confetti data [20] were the same, except the fixed modification on cysteine was set to N-ethylmaleimide (this decision was motivated by previous analysis of modifications with AA_stat [31,32]. Also, for each protease, its corresponding rule was specified, and the number of allowed miscleavages was set as follows: two for ArgC, four for AspN, six for chymotrypsin, four for GluC, two for LysC, two for trypsin. For elastase, non-specific cleavage was used. Own data from the HEK-293 cell line were searched against the proteogenomic database described in the previous work [24] for the analysis of SAPs and against the RefSeq database for the analysis of alternative splicing. Search parameters were the same as above, with fixed cysteine carbamidomethylation and variable methionine oxidation, four miscleavages allowed for GluC and two for LysC and trypsin.

**Table 1.** Group-specific false discovery rates calculated for properly cleaved tryptic peptides, miscleaved peptides with and without confirmation by their cleaved counterparts. Selected sets of shotgun proteomic data were publicly available and generated by high-resolution LC-MS/MS using Orbitrap detectors in DDA mode.

| Source of biomaterial, digestion, PRIDE accession, reference | Properly cleaved peptides |  | Single miscleavages: Peptides with 1 miscleavage site, not accompanied by cleaved peptides |  | Pairs of miscleaved/cleaved: Peptides with 1 miscleavage site, accompanied by at least 1 cleaved counterparts (coverage 2X) |  |
| --- | --- | --- | --- | --- | --- | --- |
|  | Amount | FDR, % (standard deviation*) | Amount | FDR, % (standard deviation) | Amount | FDR, % (standard deviation) |
| Human melanoma cell line 82, trypsin PXD007662 [29] | 31293 | 1.02 (0.20) | 3799 | 1.13 (0.63) | 1658 | 0.12 (0.43) |
| Human cancer cell lines, trypsin/LysC, PXD007686 [38] | 42733 | 0.92 (0.17) | 5027 | 2.08 (0.75) | 3617 | 0.06 (0.2) |
| Murine microglia culture, trypsin/LysC, PXD14466 [37] | 41685 | 0.87 (0.16) | 6474 | 2.45 (0.7) | 5680 | 0.00 (0.09) |
| Fruit fly brain, trypsin, PXD009590 [39] | 20582 | 0.94 (0.35) | 2840 | 1.33 (1.16) | 1412 | 0.00 (0.6) |
| Murine brain, fractionated, trypsin/LysC, PXD001250 [40] | 77363 | 1.00 (0.13) | 7817 | 1.53 (0.49) | 4903 | 0.04 (0.14) |

\* The standard deviation for FDR is calculated as described earlier [41].

HEK-293 proteomics data have been deposited to the ProteomeXchange Consortium via the MassIVE (https://massive.ucsd.edu) partner repository with the dataset identifier PXD030226.

### 2.6. MS1 only search to identity variants in HEK-293 cells

HEK-293 data were additionally analyzed using DirectMS1 (ms1searchpy v. 2.1.7) approach [33]. Search parameters were default except missed cleavages were set to 0, 0 and 2 for trypsin, LysC and GluC, respectively. For the analysis, only variant proteins were considered; of those, we only kept the proteins which contain at least one theoretical variant peptide in the 7-30 length range for all three proteases. Thus, only 308 variant proteins were considered in DirectMS1 results.

## 3. Results and discussion

### 3.1. Bad and good of the protein miscleavage

The most widely used protease for proteome-wide digestion in shotgun proteomics, and obviously the best in terms of specificity, is trypsin. As all enzymes, this protease has its difficulties even in nominally specific sites, which are dictated by the surrounding protein sequence context [19,34]. In the field, there is a trend to prevent the protein miscleavage by trypsin, as miscleaved peptides are less suitable for mass-spectrometry pipelines due to being longer; also, their presence inflates the search space, increasing FDR [35]. Lys-C, a protease with a similar specificity, is added to trypsin for better digestion of problematic cleavage sites [36].

When we aimed to identify and validate tryptic peptides which contained single amino acid variants originating from genomic variants [24,29] or RNA editing [8], some of those variants were found in peptides with missed cleavages. Manual inspection of the corresponding mass spectra, which was conventionally used to confirm true identifications, led us to the observation that peptide hits represented by miscleaved peptides alone were enriched with false positive results. In contrast, if the sequence variants were represented by both properly cleaved and miscleaved peptides, these results were always true [8]. Indeed, two different peptides containing the same residue of interest may be considered as independent events with multiplied probabilities. Thus, if a miscleaved peptide was identified along with one or more of its cleaved counterparts, the overlapping sequence was validated with higher probability than for any peptide alone. We then used the target-decoy analysis to illustrate this concept in selected available datasets for shotgun proteomes produced in DDA mode.

Group-specific FDR values were calculated separately, for groups of (i) properly cleaved tryptic peptides; (ii) peptides containing a single miscleavage site and lacking a confirmation with an overlapping properly cleaved peptide; and (iii) peptides containing a single miscleavage site and also confirmed by at least one overlapping properly cleaved peptide, in most cases forming pairs (Table 1). Corresponding decoy subsets were used for FDR calculations. Other groups, such as peptides containing more than one miscleavage, were also considered but not reported here as the peptides were found in low numbers. As expected, FDR values for the group of miscleaved/cleaved pairs were approximately an order of magnitude lower than for properly cleaved peptides without additional confirmation by overlapping peptides with miscleavage, with zero FDR in some datasets (Table 1). In other words, miscleaved peptides, in some cases, provide more reliable identification for the strands of amino acid sequence which have a double coverage. Interestingly, in the dataset of murine microglia [37], the proportion of miscleaved peptides is higher than in other exemplary datasets (14% of pairs vs. 5-8%, correspondingly), more likely, due to some sample preparation features. At the same time, the double sequence coverage, according to our teaching, in this dataset is also higher. As mentioned above, it is generally difficult to control trypsinolysis to guarantee efficiency. An easier and better established way to produce overlapping peptides proteome-wide is multiprotease cleavage [20].

### 3.2. Sequence coverage in multiprotease proteomic datasets

The concept of sequence coverage by multiple reads as defined here relates mostly to more reliable identification of single amino acid or peptide events in proteome, such as amino acid sequence variants, splice junctions etc. In this case, the coverage can be defined as easy as the number of distinct peptides spanning the site of interest, with the majority of conventionally identified sites having a coverage of 1X. At the same time, to compare between datasets and methods for retrieval of the coverage it is convenient to estimate its fold proteome-wide. This task is more complicated because a whole proteome, similarly to the genome, contains many degenerated sequences which may corrupt the coverage estimation. Indeed, there is no linear “proteome” onto which all peptides can be mapped; instead, there are concurrent isoforms, splice products, etc. One way of approaching this issue is to map all peptides onto the genome directly; however, in this work we take a simpler approach and consider a subset of identified peptides that can be mapped to proteins without ambiguity.

To calculate the overall proteome coverage, we started with the output of Scavager postsearch validation tool [28] (Fig.1). Scavager produces several tables with identified PSMs, peptides, proteins and protein groups. Each of the tables is filtered to the desired FDR level (1% in our case). Each of the protein groups has a “leader” protein, and it is known that no proteins in any group except the leader have any unique peptides. We considered all identified peptides, and considered which proteins they come from. Then, we only kept the peptides which have exactly one protein group leader in their list of proteins; these peptides could be unambiguously mapped to this leader protein. The leader proteins from this peptide subset formed the simplified, “linear” model of the proteome, for which the average coverage can be easily calculated. This approach to define the linear proteome was similar to the method of genomic SNP calling where the sequences shared between different genes are excluded from analysis to provide so-called “mappability” [42].

**Figure 1.**
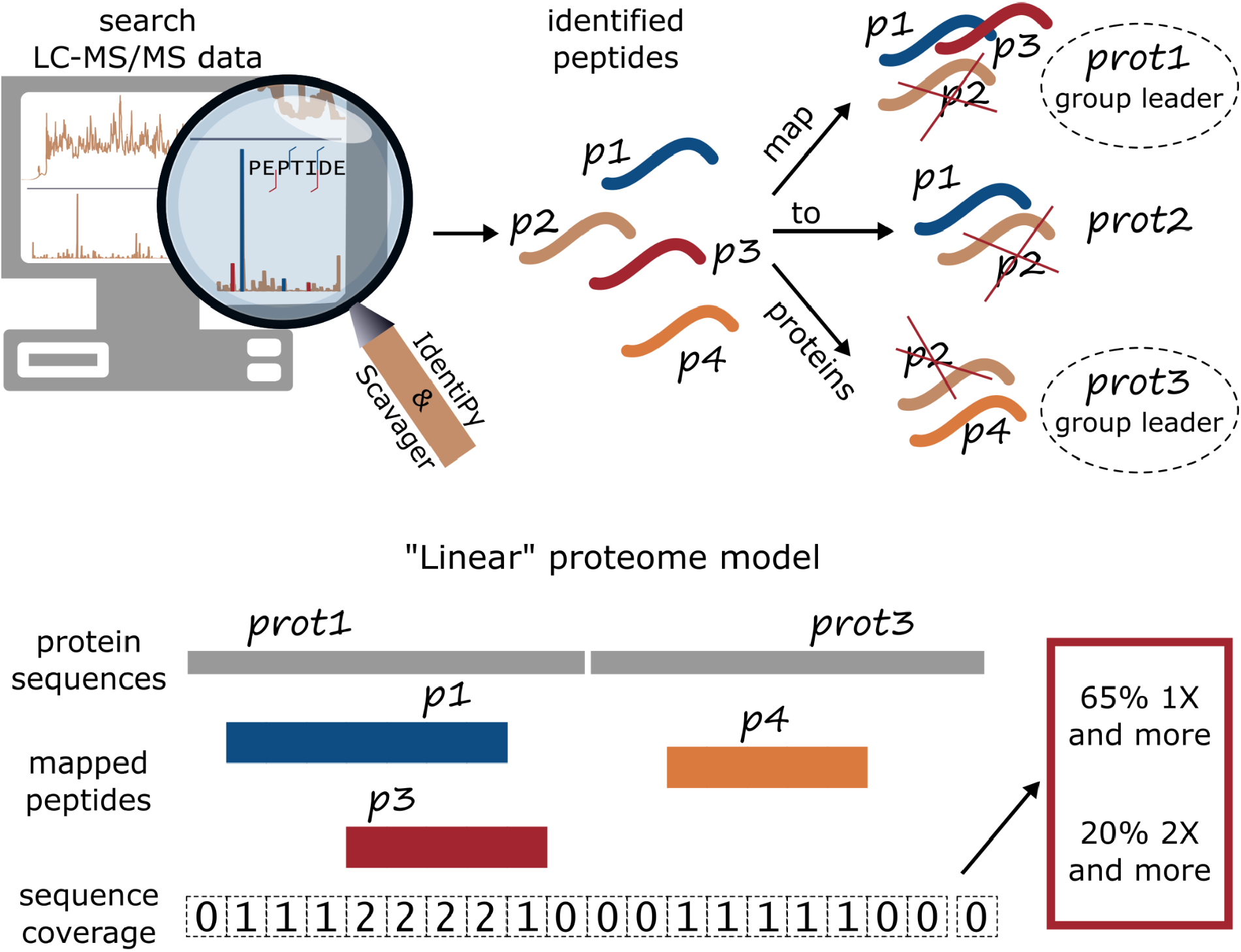
The concept of proteome-wide sequence coverage by multiple reads. To quantify the multiplicity of sequence coverage throughout the observed proteome, we build its “linear” model, composed of “group leader” proteins as reported by Scavager tool [28]. Peptides shared between protein groups are discarded, and the rest are mapped onto the “linear proteome”. The “linear proteome” is represented computationally by an array of integer values (initialized with zeros), one for each amino acid position in every detected “group leader” protein. Then, each mapped peptide increments the values corresponding to its place in the protein sequence. Afterwards, the distribution of array values is summarized as proteome coverage statistics.

To calculate the proteome coverage, each of the leader proteins is represented with an array of zeros; each position in the array corresponds to a single amino acid residue in the protein sequence. For each of the distinct peptides from the subset defined above, the corresponding portion of the protein array is incremented by one. After that, all protein arrays are concatenated, and the distribution of values in the resulting array is considered as a representation of proteome coverage. Note that only unique peptides were considered in this procedure; repetitive PSMs for the same peptide were not taken into account, even when they come from samples treated with different cleavage agents. This is done to exclude possible systematic errors that could lead to repetitive incorrect peptide assignments.

Two most frequent enzymes used for shotgun proteomics are trypsin and LysC, which have similar specificity, except that digestion after arginine residues is not feasible by LysC. However, in most cases these enzymes are used concurrently. For purposes of this work, it was interesting to find data where two enzymes acted separately, to estimate the gain in coverage due to the difference in their specificities. A good example of such data was a human brain dataset by Wingo et al [30], where trypsin and LysC were used separately. This proteome, deeply covered using prefractionation of enzymatic lysates, was characterized by 8598 protein groups in total, according to our reprocessing of the raw data. First, we estimated if miscleaved peptides add to the sequence coverage in the trypsin subset of this brain proteome. At a total proteome coverage of 24%, about two percent of the linear proteome sequence was covered 2-fold by miscleaved peptides (see Section 3.1). Combining trypsin data with LysC runs slightly increased the general coverage to 28%. At the same time, coverage of 2-fold or more reached about 6%. Thus, about 22% of all identified proteome regions in this dataset were covered by multiple reads, when both trypsin- and LysC-digested samples were considered.

A unique collection of multiprotease shotgun proteomic data is Confetti, which represents a multiprotease proteome map of the HeLa cell line [20]. Expectedly, the use of seven proteases with different specificities enhanced the multiple coverage up to 35% of the visible proteome regions, while the overall proteome coverage was relatively low (20%) due to relatively poor sensitivity, with 4,8 thousand proteins identified (Table 2).

**Table 2.**
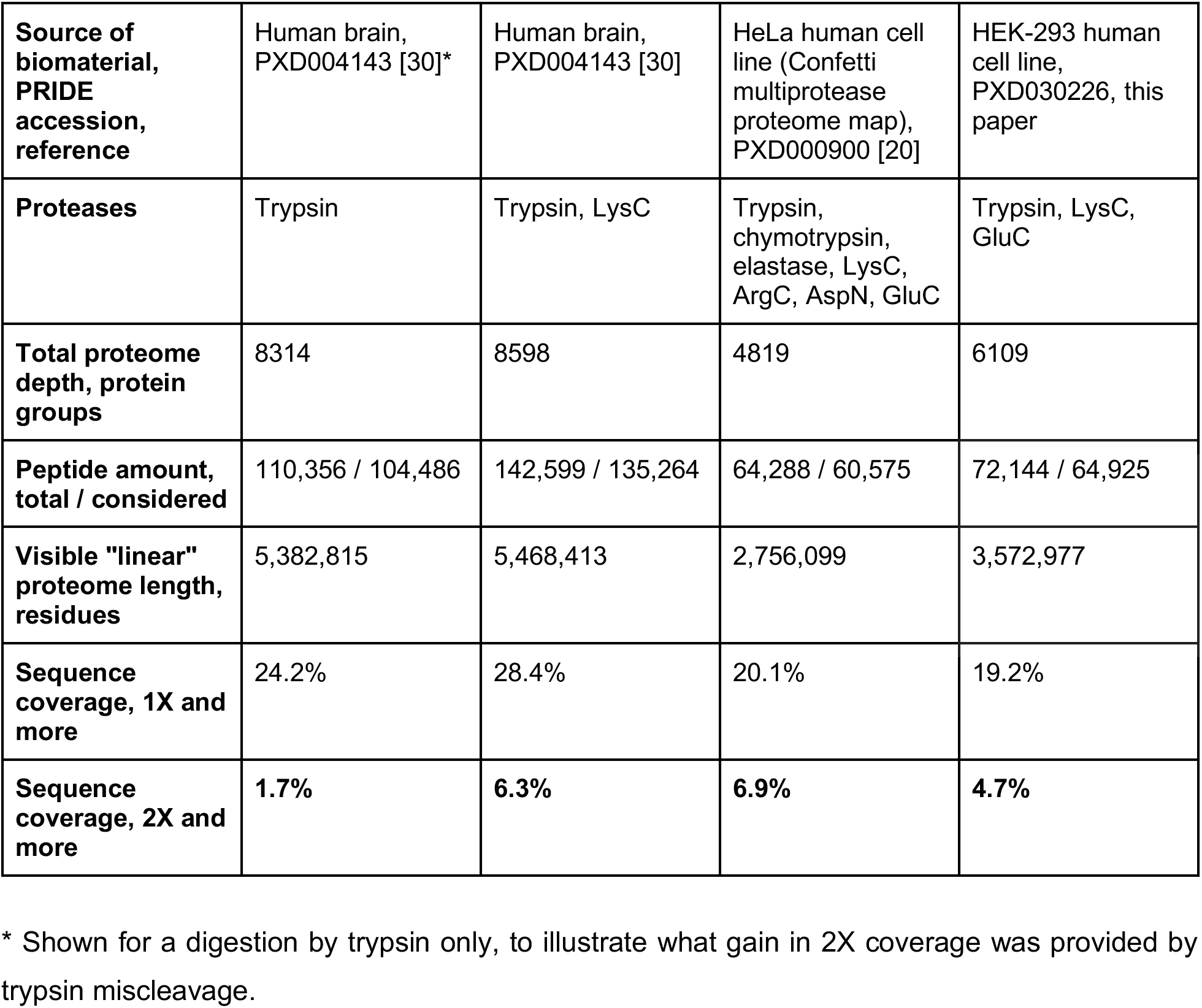
Characteristics of the sequence coverage calculated proteome-wide for shotgun proteomic datasets generated using multiple proteases

|  |  |  |  |  |
| --- | --- | --- | --- | --- |
| **Source of biomaterial, PRIDE accession, reference** | Human brain, PXD004143 [30]* | Human brain, PXD004143 [30] | HeLa human cell line (Confetti multiprotease proteome map), PXD000900 [20] | HEK-293 human cell line, PXD030226, this paper |
| **Proteases** | Trypsin | Trypsin, LysC | Trypsin, chymotrypsin, elastase, LysC, ArgC, AspN, GluC | Trypsin, LysC, GluC |
| **Total proteome depth, protein groups** | 8314 | 8598 | 4819 | 6109 |
| **Peptide amount, total / considered** | 110,356 / 104,486 | 142,599 / 135,264 | 64,288 / 60,575 | 72,144 / 64,925 |
| **Visible "linear" proteome length, residues** | 5,382,815 | 5,468,413 | 2,756,099 | 3,572,977 |
| **Sequence coverage, 1X and more** | 24.2% | 28.4% | 20.1% | 19.2% |
| **Sequence coverage, 2X and more** | 1.7% | 6.3% | 6.9% | 4.7% |

\* Shown for a digestion by trypsin only, to illustrate what gain in 2X coverage was provided by trypsin miscleavage.

To demonstrate the sequence coverage approach with our own data, we used a multiprotease digestion scheme on the HEK-293 model cell line which was previously studied in our work focused on its amino acid variants [24]. Proteins extracted from the same sample used before were digested separately by three proteases, including trypsin, LysC and endoproteinase GluC, followed by shotgun analysis of the digests in DDA mode in three technical replicates. These analyses yielded 5435, 4950, and 3307 protein groups for trypsin, LysC, and GluC, respectively. Combined, 6109 protein groups were identified. These data provided a general coverage of the linear proteome about 19%, which is lower than for the brain proteome data described above [30]. Note that the brain proteome results were obtained using prefractionation of the peptide mixture. At the same time, a relative proteome coverage of 2X and more for our HEK-293 data was higher than in the brain data and reached about 25% of the covered regions vs. 22% in the brain dataset. This relative improvement, indicated as significant by Fisher’s exact test, was provided by the addition of GluC to the protease set. At the same time, much deeper analysis in the brain dataset provided a two-fold gain in absolute characteristics of multiple coverage, i.e. about 0.34 mln amino acid residues covered 2x and more vs. 0.17 mln in the HEK-293 data. A high portion of identified proteome with multiple coverage in the Confetti dataset (35%) also looks not so impressive in absolute numbers spanning 0.19 mln residues (Fig.2A).

**Figure 2.**
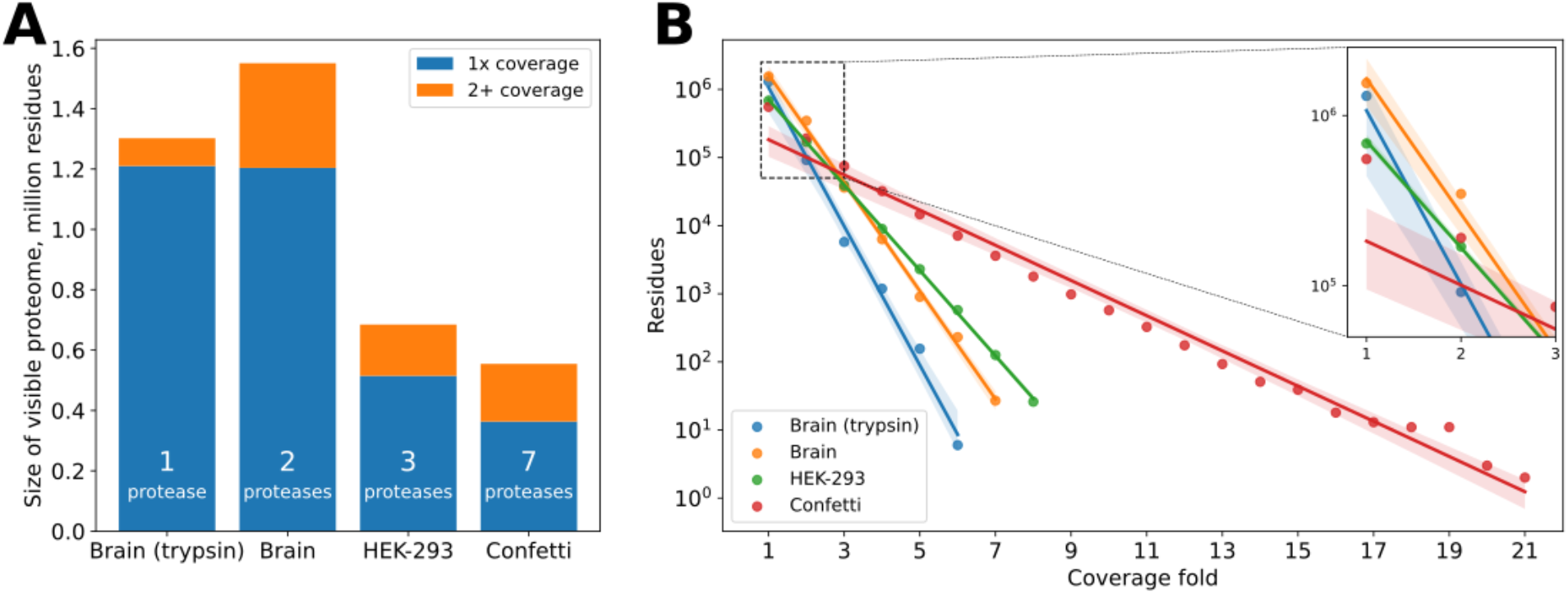
Size comparison of visible proteome sizes in the human brain, HEK-293 and Confetti datasets. The brain sample is sampled much deeper due to fractionation; addition of LysC analysis adds more coverage to the same sequence regions in already detected proteins. Increasing the number of proteases used in parallel results in a higher proportion of multiply-covered sequences (*A*) Distributions of coverage multiplicity (fold) values in detected proteins follow exponential trends (straight lines in logarithmic scale). Line parameters depend on the number of proteases (slope) and sensitivity in a single analysis (intercept) (B).

Not many datasets with distinct treatment of multiple proteases are publicly available. However, the three examples provided here confirmed an intuitively clear conclusion that a gain in proteome sequence coverage in multiple reads may be reached by two experimental conditions. First, a number of proteases used for independent, simultaneous proteome analysis may be increased. Second, both the general coverage and the coverage by multiple reads may be increased with deeper proteomic analysis, attainable e.g. by prefractionation of the peptide mixture and analyzing the fractions separately. There are many examples of successful use of multiprotease analysis to increase the proteome coverage [20–23]. The novelty of our approach is the use of this analysis to confirm selected single amino acid events in contrast to other works which were focused on more complete inference of whole proteins.

As shown in Fig.2B, the total length of the proteome covered with at least *k* distinct peptides decreases exponentially as a function of *k*. This is expected if we consider that these peptide identifications are largely independent. The relationship becomes linear when switching to the logarithmic scale. Straight lines obtained from least-squares linear fit succinctly describe the multi-proteases analysis: the slope of the line depends on the number of proteases used (the more proteases, the flatter), while the intercept corresponds to the depth of the proteome analysis of each of the parallel analyses. For example, the brain datasets processed with trypsin only and with two proteases have nearly identical total numbers of residues covered exactly once (two left blue bars in Fig.2A, left-most blue and yellow points in Fig. 2B, see inset). Thus, the blue and yellow lines have the same value of intercept, however, the addition of LysC results in a flatter slope.

### 3.3. Multiprotease shotgun analysis of HEK-293 proteome: sequence coverage of single amino acid variants

A modified genomic database for peptide identification with added amino acid variants was taken from our previous study [24]. The new dataset made it possible to estimate (i) the reproducibility of amino acid variant identification between different datasets; (ii) the gain in variant identification from the addition of treatment by GluC, a protease with complementary specificity to trypsin and LysC; and, finally, (iii) the portion of coverage of these variants by multiple peptide reads.

Fortunately, the majority of 36 protein sequence variants found in HEK-293 cells (Table 3) were also identified before in at least one of three reported proteomes of this cell line [24,43,44] described in detail in our background paper [24]. More specifically, 28 of them were reproduced here (Table 3). This amount of variants was generally comparable with two background studies, where 32 and 38 variants were reported from our own [24] and Geiger et al. data [44]. Reprocessing the data for the much deeper proteome by Chick et al [43] yielded 84 amino acid variants identified. The use of GluC in our study led to the addition of 5 new variants which fell into parts of sequences poorly compatible with mass-spectrometric identification after digestion by trypsin/LysC. Two variants covered only by GluC-generated peptides in our data, namely, in PIH1D1 and TP53BP1 proteins, were also detected by tryptic peptides in the data by Geiger et al [44] and by Chick et al [43], respectively (Table 3).

**Table 3.**
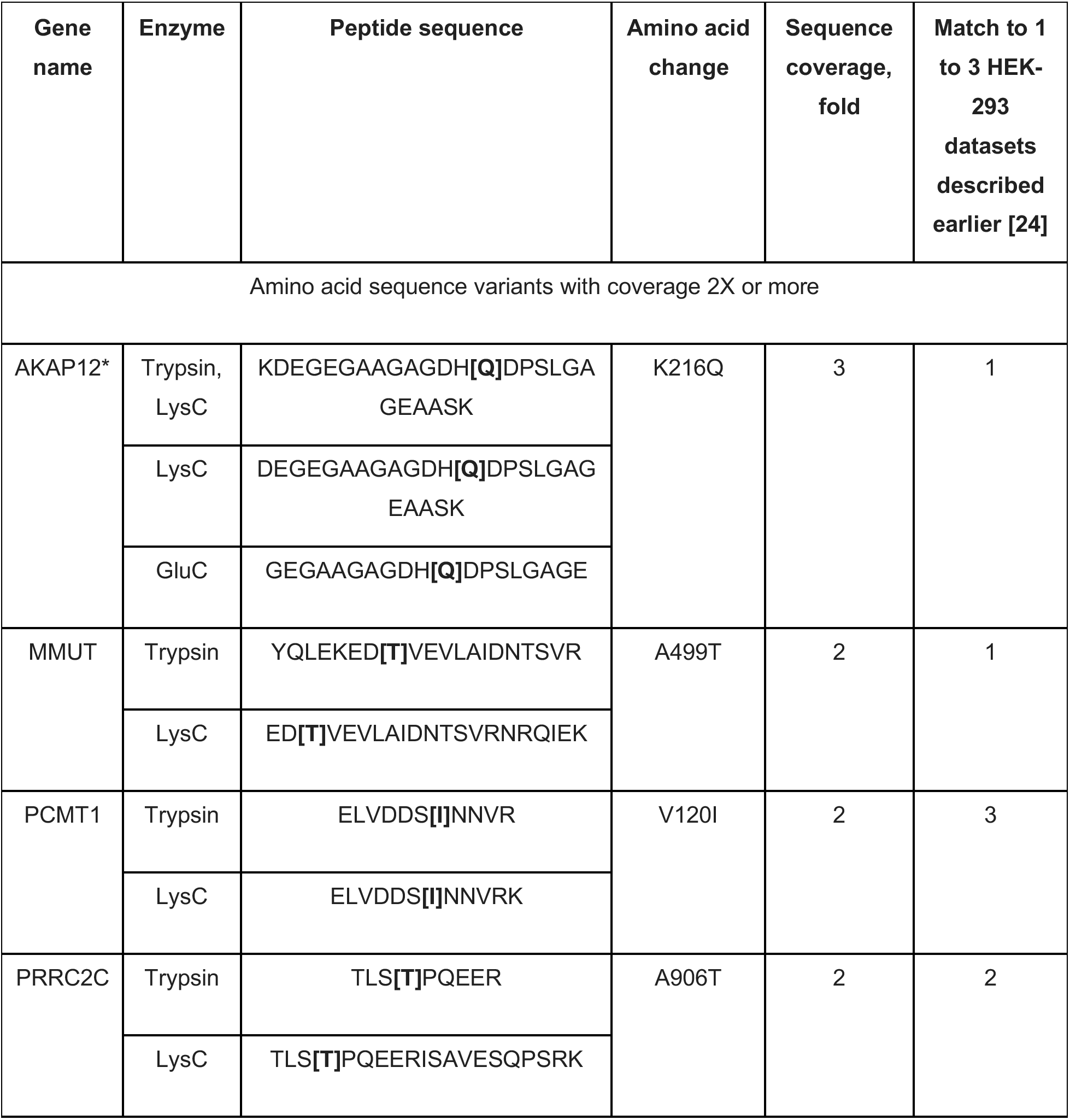
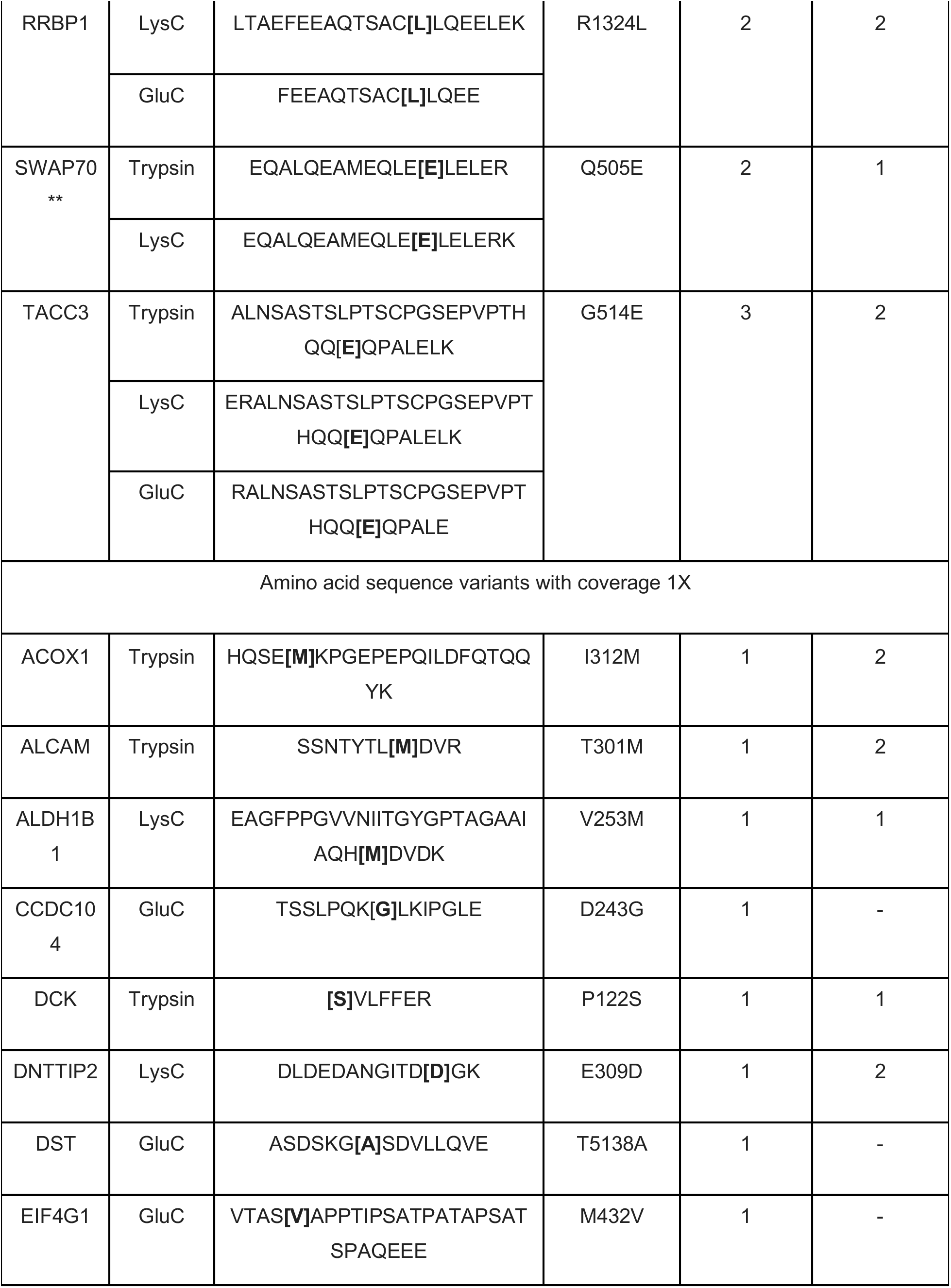
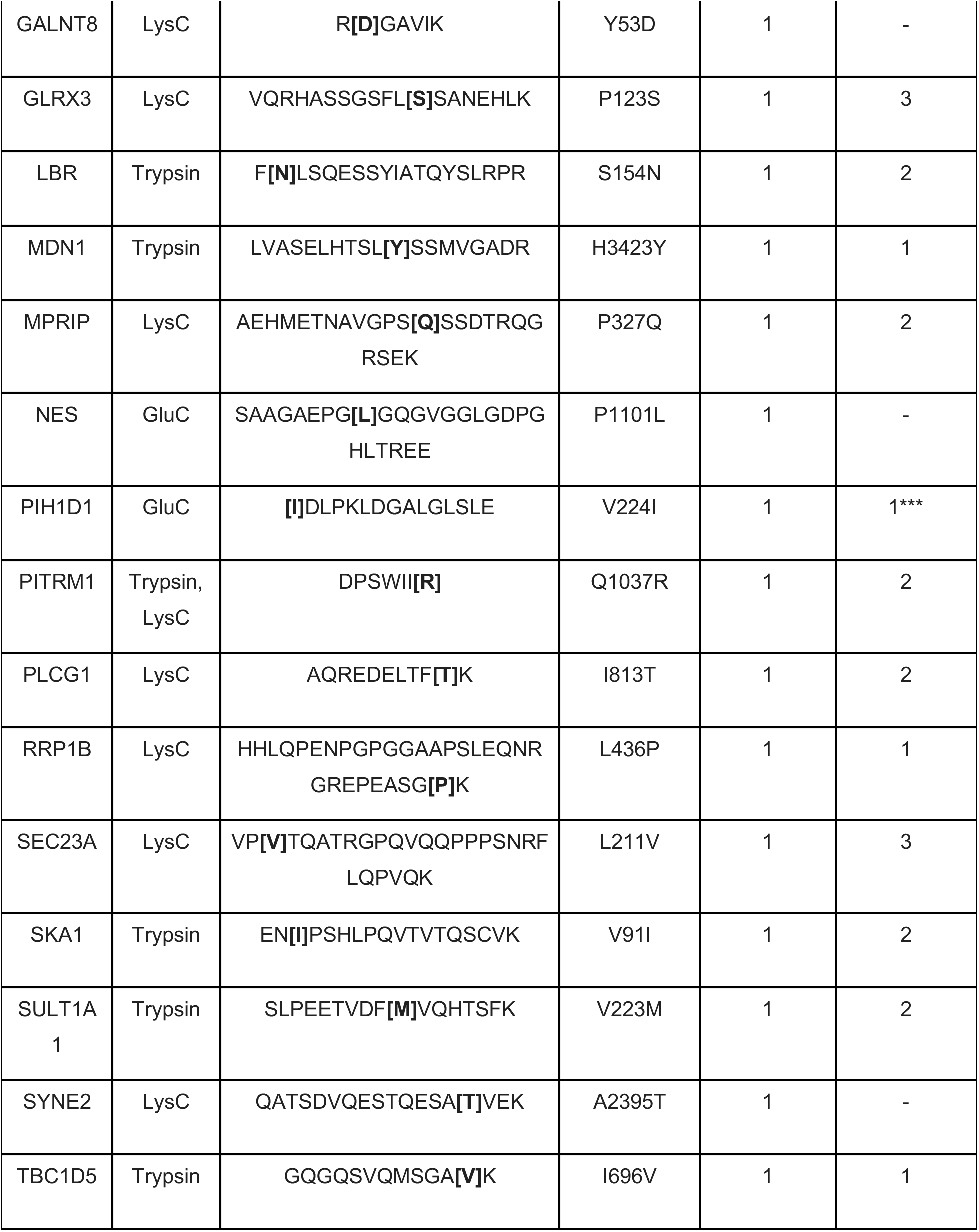
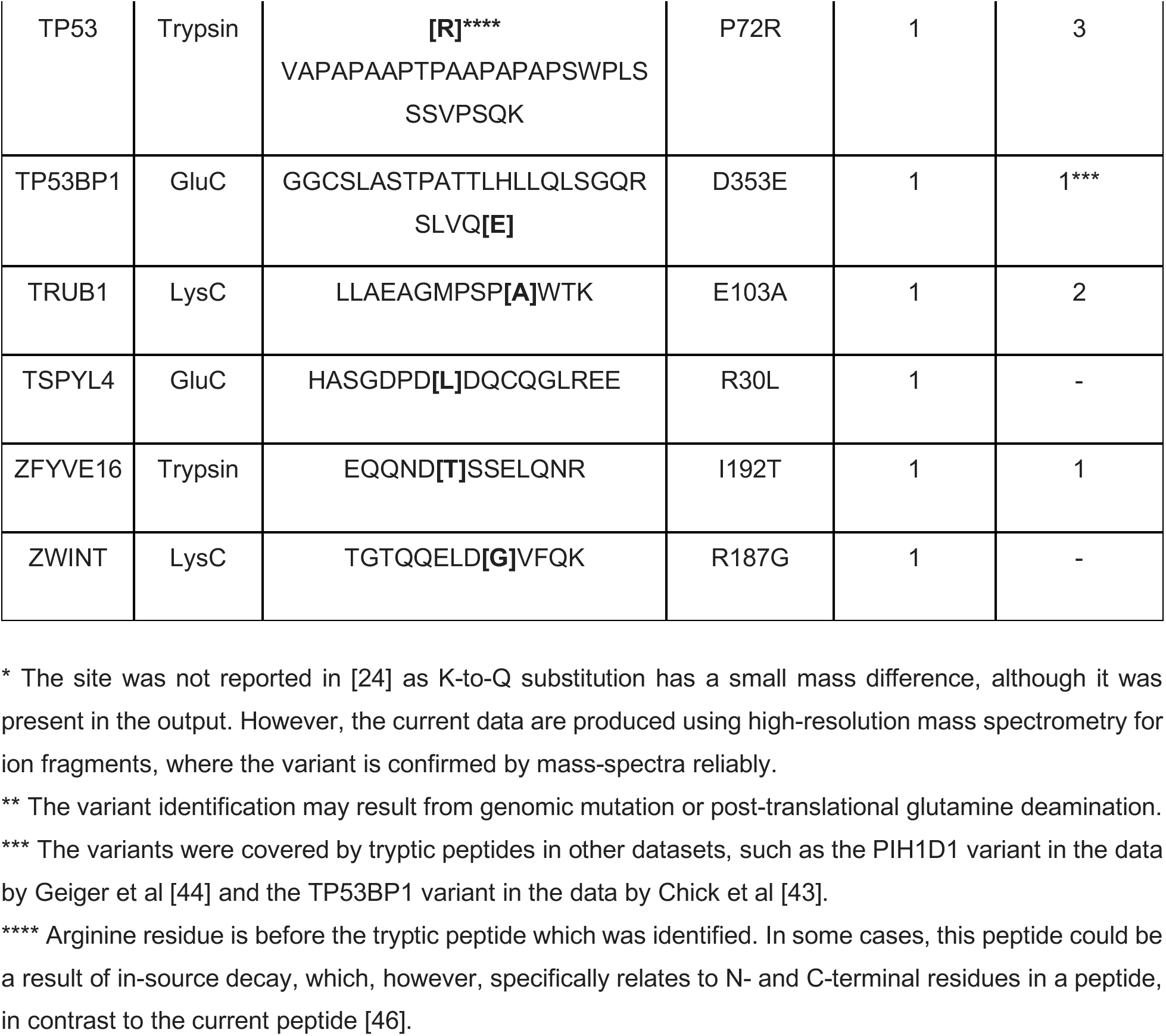
Missense genomic variants identified in the shotgun proteome of the HEK-293 cell line using the multi-enzyme approach to provide enhanced sequence coverage by distinct peptide reads. Genomic database supplemented by amino acid variants was taken from the previous study [24]. The amino acid numbering is represented according to the isoform 1 in the NextProt knowledgebase, data release: 2021-02-15 [45].

The variant sequence coverage was estimated as exemplified in Fig.3. The portion of variants with coverage 2X or more was quite similar to that observed proteome-wide. Namely, of 36 variants identified with use of any enzyme, seven were covered by at least two distinct peptides (19.4%). Of these, the K216Q sequence variant of A-Kinase Anchoring Protein 12 (AKAP12) and the G514E variant of cancer-related Transforming Acidic Coiled-Coil Containing Protein 3 (TACC3) were covered by three distinct peptides generated by all three enzymes used, i.e. these sites had 3X coverage. Variants of proteins encoded by MMUT, PCMT1, PRRC2C, and SWAP70 all had 2X coverage provided by trypsin and LysC. The identification of the Q505E variant of SWAP70 may either result from genomic mutation or post-translational glutamine deamination [6]. However, we report it for illustration of our approach. Finally, the R1324L variant of RRBP1 was also covered 2-fold by LysC and GluC (Table 3).

**Figure 3.**
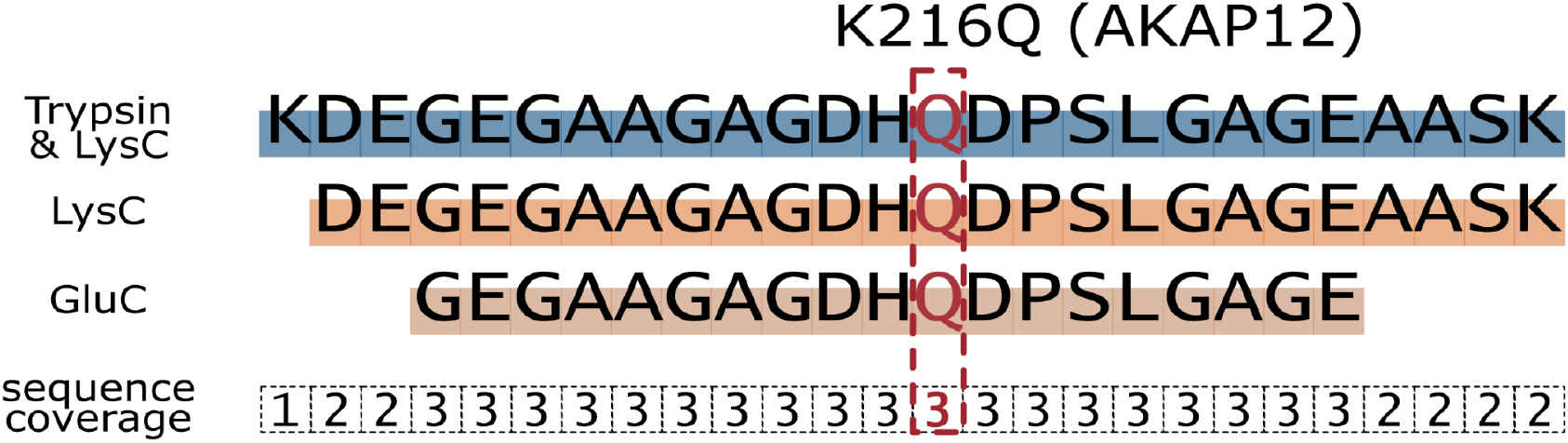
Sequence coverage of the K216Q single amino acid variant of A-Kinase Anchoring Protein 12 (AKAP12) by three distinct peptides generated by trypsin, LysC and GluC.

One of the ways to increase reliability of findings confirmed by specific PSMs, such as single amino acid variants discussed above, is inspection of mass spectra. Mass spectra of the variants with the sequence coverage of two or more (Table 3) were visualized by xiSPEC spectrum viewer [47] to determine if the specific amino acid residue at the alleged mutation site was confirmed by the corresponding b- and/or y-ions. For the K216Q variant in AKAP12, the corresponding glutamine residue was confirmed by fragmentation mass-spectra attributed to all three distinct peptides containing the variant (Fig. 4). Correspondingly, single amino acid variants in PCMT1, PRRC2C, and RRBP1 were confirmed by fragment mass spectra for all distinct peptides containing these variants (Figs. S1-S3, respectively). Further, the G514E variant in TACC3 was supported by fragment ions in two of three possible cases (Fig.S4). The variants in MMUT and SWAP70 were confirmed by fragmentation of one of two peptides (Figs. S5-S6).

**Figure 4.**
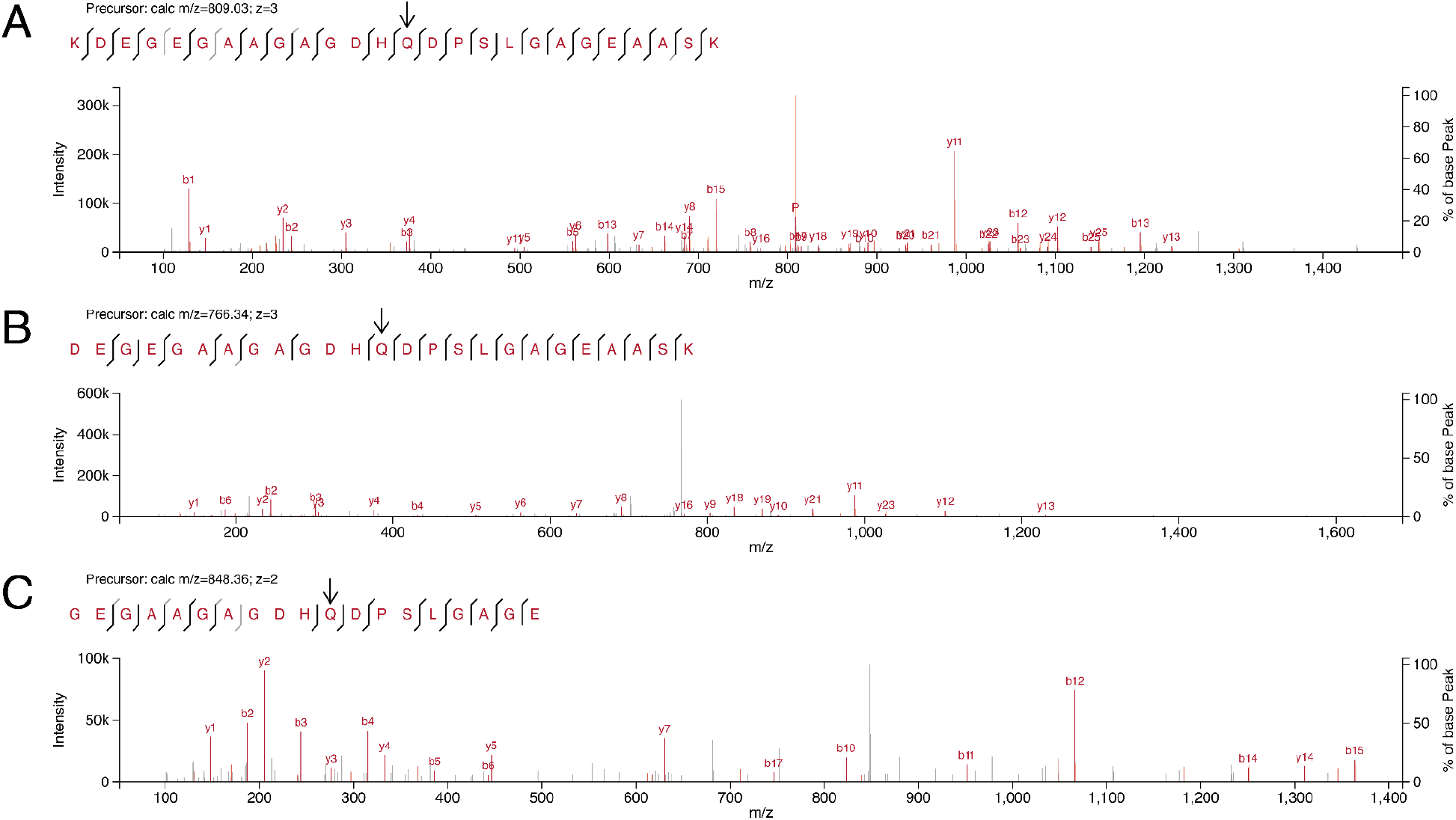
Fragmentation mass spectra of peptides of A-Kinase Anchoring Protein 12 (AKAP12) containing a genetically encoded K216Q amino acid sequence variant. Mass spectra recorded from the shotgun proteomes of HEK-293 cell line after protein digestion by trypsin (A), LysC (B), and GluC (C). Vertical bars around amino acid letters show that residues were confirmed by y-ions (upper branches) or b-ions (under branches). Mass spectra were visualized by xiSPEC spectrum viewer [47].

A question might arise whether the proteins with variants covered 2X or more were highly abundant in HEK-293 proteome. Using NSAF label-free quantitation method [48], percentiles of abundance could be defined for each of them in the trypsin-digested dataset. Indeed, AKAP12, a structural component of protein kinase A signalosome, was abundant in this cell line, placed at the 84th percentile. However, the other six proteins on the list were more or less evenly distributed along the list, for example, with 28th percentile for SWAP70. Thus, the coverage of specific sequences is controlled by physical properties of the corresponding peptides.

### 3.4. Sequence coverage of splice junctions for alternative splicing

Alternative splicing site coverage was calculated in the following way. Based on the RefSeq reference genome annotation (GRCh38) all consecutive coding sequence (CDS) pairs were considered in all annotated transcripts of all human genes. Theoretical peptide sequences were generated by translating these pairs and performing *in silico* digestion in accordance with enzyme specificity of trypsin, LysC, or GluC, allowing for 2, 2 and 4 miscleavages, respectively. Peptides lying entirely within a single CDS, as well as peptides present in all protein products of the given gene, were discarded. Thus, only alternative splicing peptide products were kept for each possible site.

For 116,018 possible alternative splicing events in 12,968 genes, a total of 464,078 theoretical confirming peptides were predicted for trypsin, 253,815 for LysC and 7,118 for GluC. To calculate the coverage of each splicing site, a database search was performed against the regular RefSeq protein database. Then, reliably identified peptides confirming each site were counted across all experiments.

In our experiments, peptide evidence was found for 4209 alternative splicing events in 2101 genes of HEK-293 cells (Table S1). However, some peptides could not be mapped unambiguously to a single gene due to paralogy. A total of 273 unique junction-spanning peptides from paralogous proteins were identified; they were excluded from further analysis, leaving 3893 unique peptides. Additionally, some of these peptides attributed to a single gene still did not point unambiguously to a specific alternative splicing event, due to the fact that the splicing site was too close to the start or end of the peptide sequence. In those cases, almost all of the sequence coding the peptide is located within one exon, and the remaining few nucleotides are not enough to unambiguously identify the other exon. These cases were also excluded, leaving 3865 peptides as evidence for 3350 splice events. Of these events, only 478 (or 14%) were covered 2-fold or more. This percentage is much lower than for the total “linear” proteome, where it was about 25%. This observation can be presumably explained by decreased abundances of alternatively spliced proteoforms, as well as by their enrichment by lysine and arginine sites, limiting identification by trypsin and LysC [49].

### 3.5. MS1 approach to confirm amino acid sequence variants in HEK-293 proteome

An approach of express proteomic analysis which uses only precursor *m/z* values without MS/MS, called DirectMS1, was recently elaborated by authors of this work [33]. In this method, peptide-level FDR is usually uncontrolled and high, which makes variant identification tricky. However, in the case of three different proteases, the variant could be reliably confirmed if it was detected in three distinct peptides. In our analysis, two variants met this criteria, K216Q in the abundant AKAP12 protein, which was also covered 3X in conventional analysis (see Table 3), and, unexpectedly, I813T from PLCG1. In addition to the AQREDELTFTK peptide identified from the LysC subset (Table 3), DirectMS1 identified a tryptic fragment, EDELTFTK, and a GluC-produced fragment, DELTFTKSAIIQNVE. Probably, these peptides failed to pass the intensity threshold during conventional MS/MS-based data acquisition. Indeed, relative MS1 intensities for the LysC, tryptic and GluC peptides were 8×10^6^ (the one detected in MS/MS analysis), 1×10^6^ and 4×10^5^, respectively. Thus, the orthogonal identification method, based on exact *m/z* of precursor ions, added a variant hit to the results of MS/MS search.

## 4. Concluding remarks

The omics technologies today generate increasingly large data arrays providing biologically relevant information at an accelerating pace compared with the pre-genomic era in molecular biology. However, the omics, while providing the ways to measure thousands of molecular variables simultaneously, are suffering from reliability issues compared with the one-by-one approaches of the old days. New discoveries based on the high throughput omics are not self-sufficient and need extensive validation steps by orthogonal methods. This limits wider and more useful acceptance of omics in some applications, such as the clinical ones. NGS improved significantly the reliability issue in genomics, thus, paving its way to clinical discoveries and diagnosis [50].

However, mass spectrometry-based shotgun proteomics, despite the recent progress in its performance, is still lacking reliability in identification of every single amino acid of proteins proteome-wide. Here we considered a more conservative way to interpret the proteome analysis results obtained in the context of proteogenomic studies using data-dependent acquisition mode to distill the more reliable part of them, better suitable for clinical discoveries. Instead of considering all variant peptides identified at a given FDR level as equal, we suggest their further ranking using the sequence coverage by multiple reads approach in a similar way as it is done in nucleic acid sequencing for the calling of single nucleotide variants. Multiple reads for each letter in the proteome sequence can be obtained by overlapping distinct peptides, which confirm the presence of certain amino acid residues in the overlapping stretch with much lower FDR compared with 1% accepted for the whole group of identifications. These overlapping distinct peptides can be formed by, first, the pairs of miscleaved tryptic peptides and their fully cleaved counterparts, and, second, the peptides generated by several proteases with different specificities applied to the same specimen. The corresponding digests should be analyzed separately using the multiprotease proteome analysis workflow well known in proteomics [20].

Note that contrary to transcriptomics, which provides rich sequence coverage content, the coverage of each protein by peptide “reads” is relatively small in proteomics. For example, the coverage of human proteome in exemplary datasets, even with a single read, is typically in the range of 20% to 30%, with 5% to 7% of the proteome covered two-fold or more. Similarly, of 36 single amino acid variants identified here for the HEK293 cell line, 7 were covered at least twofold. However, these 7 variants can be considered as bullet-proof identifications which can be reliably explored further in clinical studies.

The approach of sequence coverage with multiple reads may be useful for clinical applications, for example, identification of cancer missense mutations which may serve as neoantigens, including experimental schemes of neoantigen vaccine production [51]. More reliable validation of cancer mutations at the proteome level may facilitate prioritization of candidate neoantigens for personalized vaccines [52]. Further, actionable or diagnostic cancer mutations and splice junctions may be identified more reliably with the proposed method, which would make it possible to omit a further validation stage, such as targeted proteomics or antibody-based methods. Outside of medicine, sequence coverage may validate basic scientific findings derived from the identifications of short sequences, such as novel classes of micro proteins derived from RNAs which were previously thought as non-coding [53].

The work was funded by the Russian Science Foundation, grant #20-15-00072, to S.M. We thank the Center for Precision Genome Editing and Genetic Technologies for Biomedicine, Federal Research and Clinical Center of Physical-Chemical Medicine of the Federal Medical Biological Agency for providing computational resources for this project.

The authors have declared no conflict of interest.

