## Supplementary figures and images for "Validating amino acid variants in proteogenomics using sequence coverage by multiple reads"

### Fig_S1.png

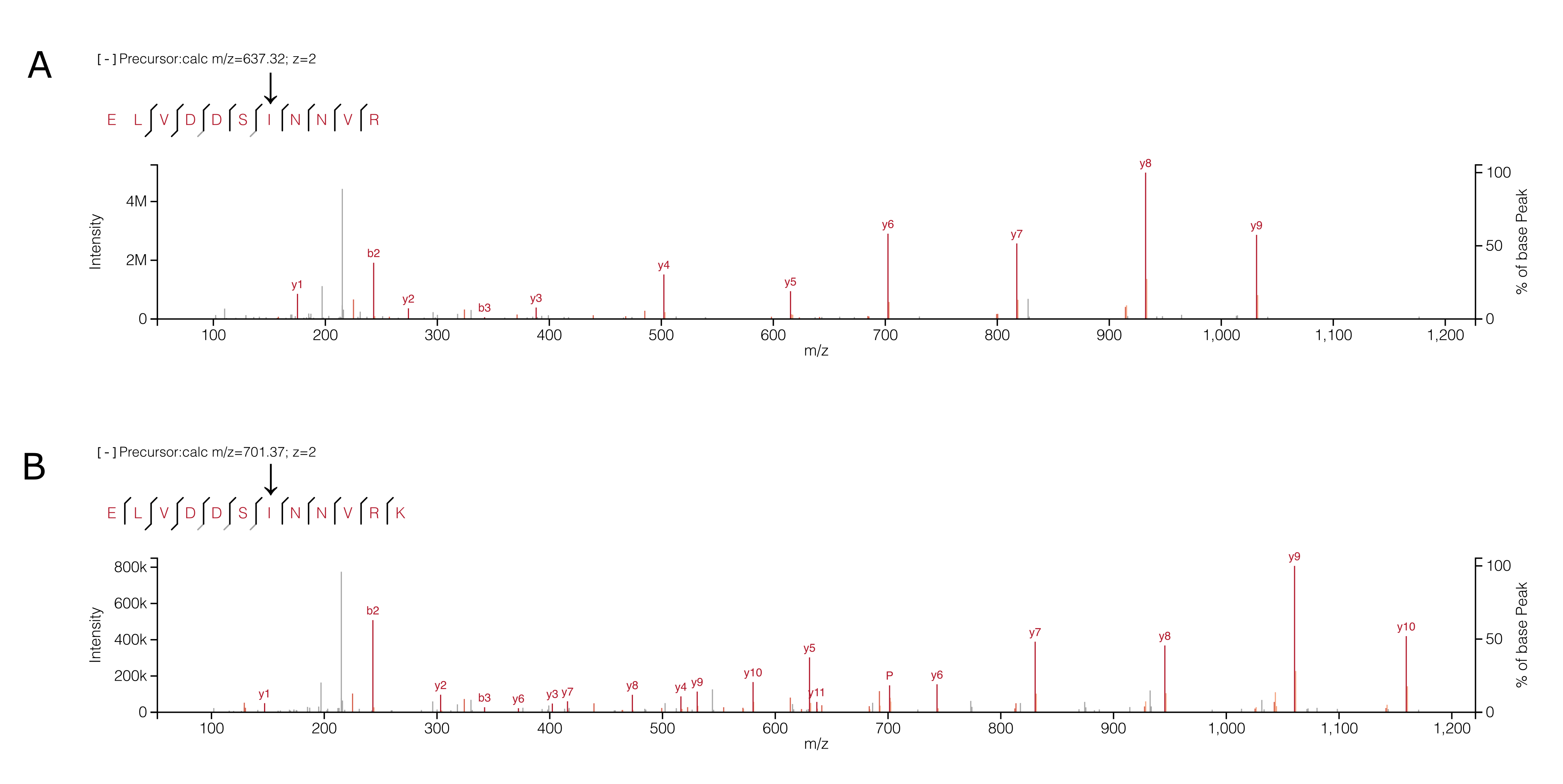

### Fig_S2.png

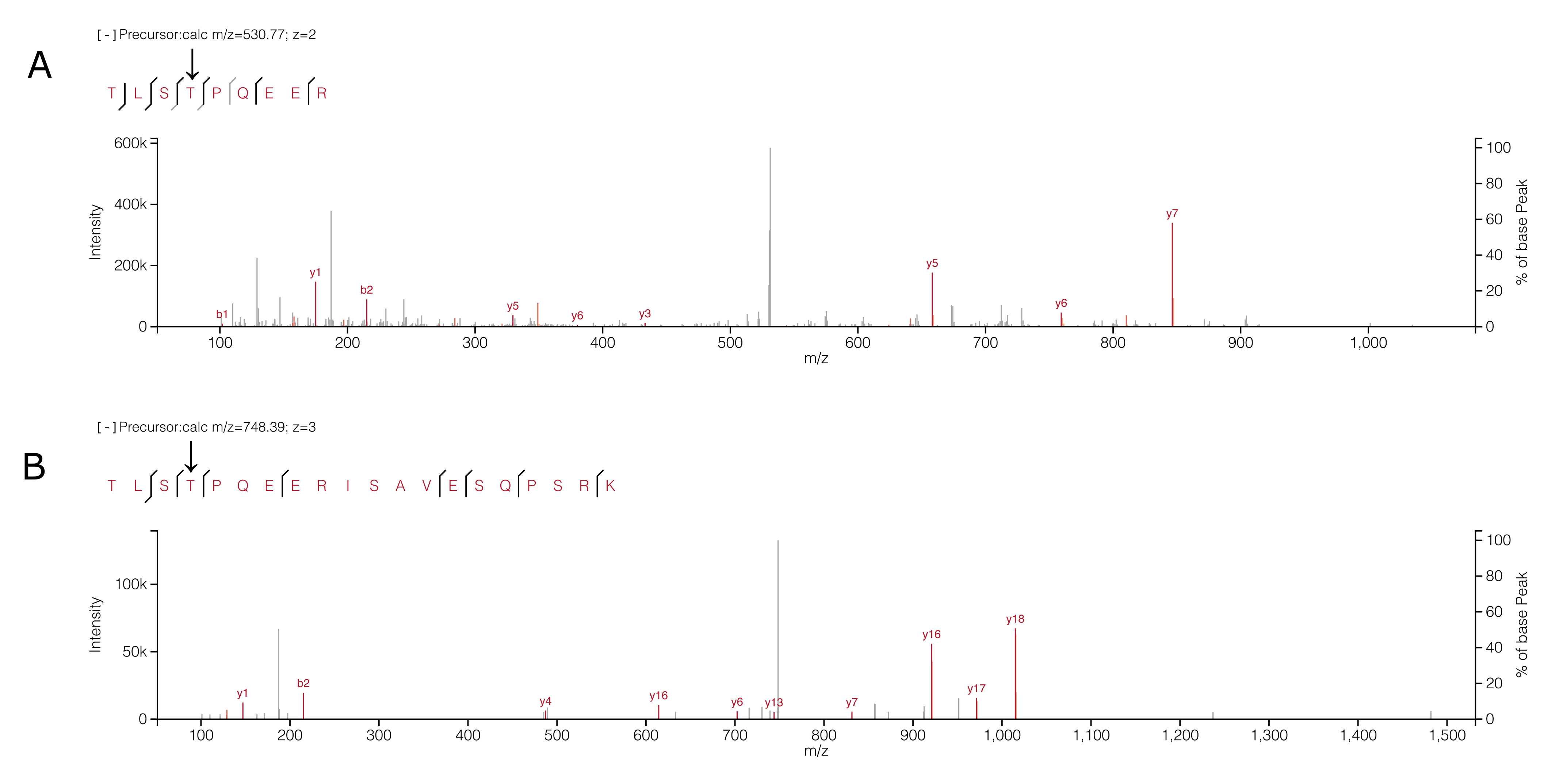

### Fig_S3.png

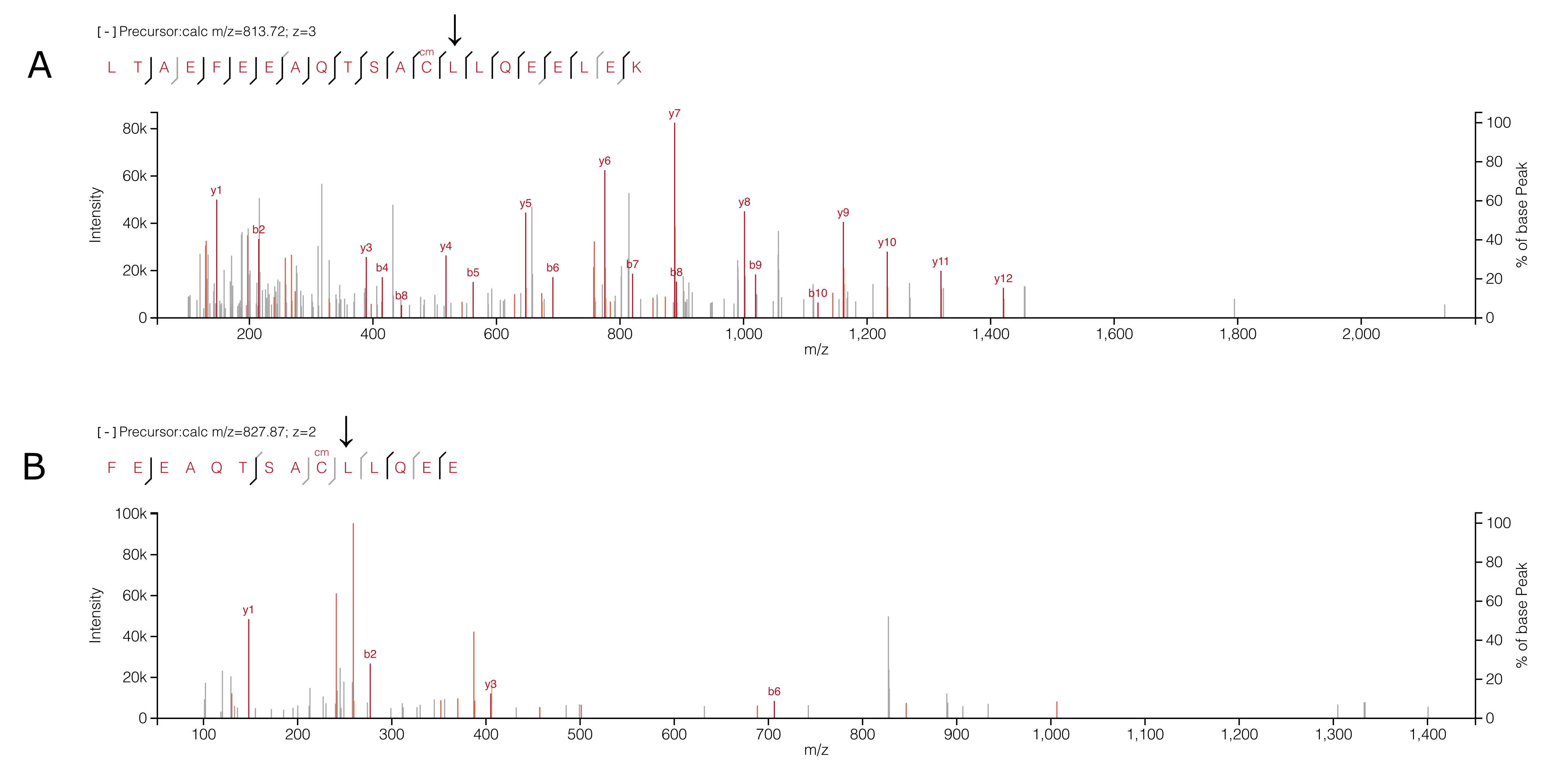

### Fig_S4.png

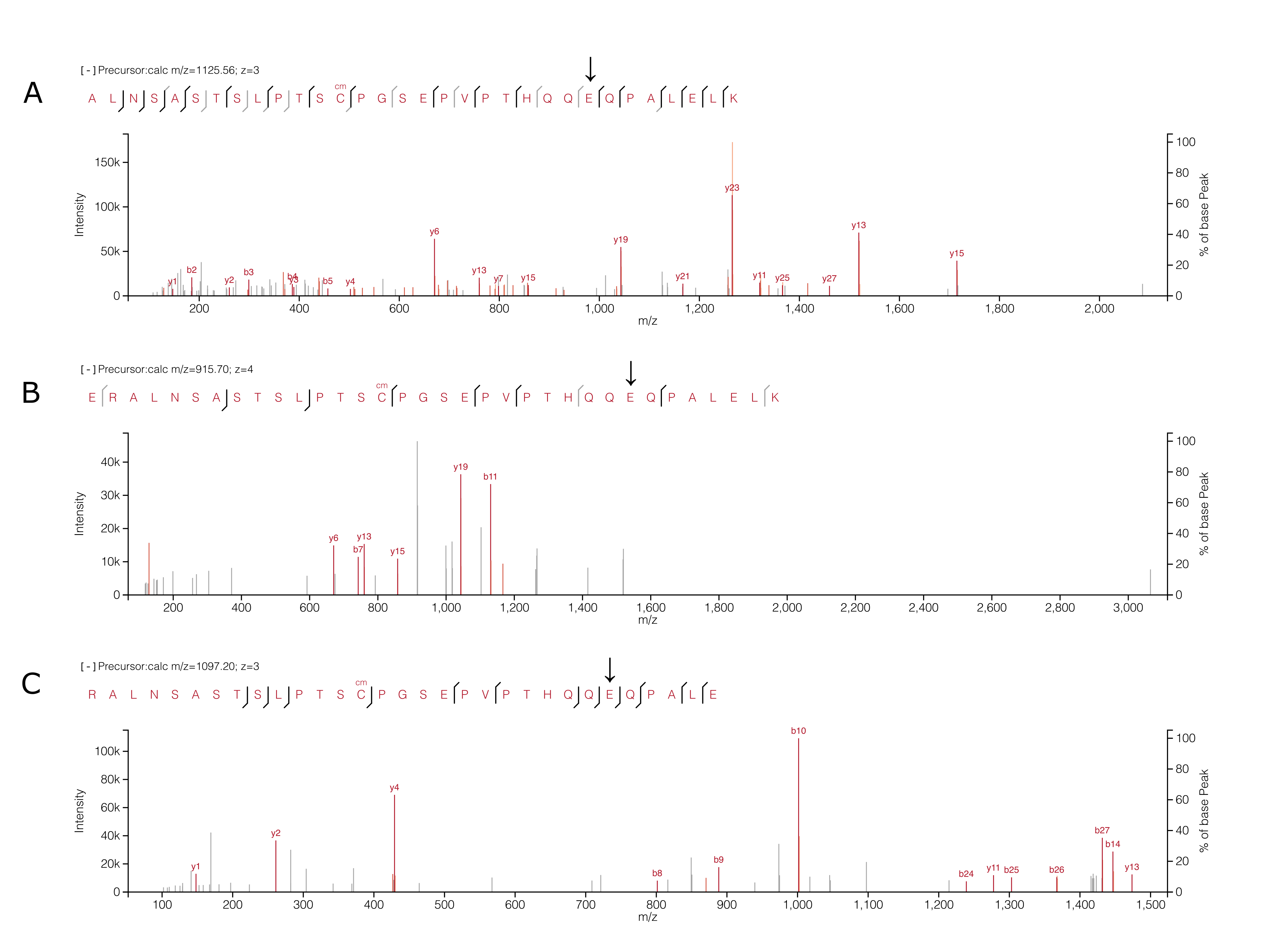

### Fig_S5.png

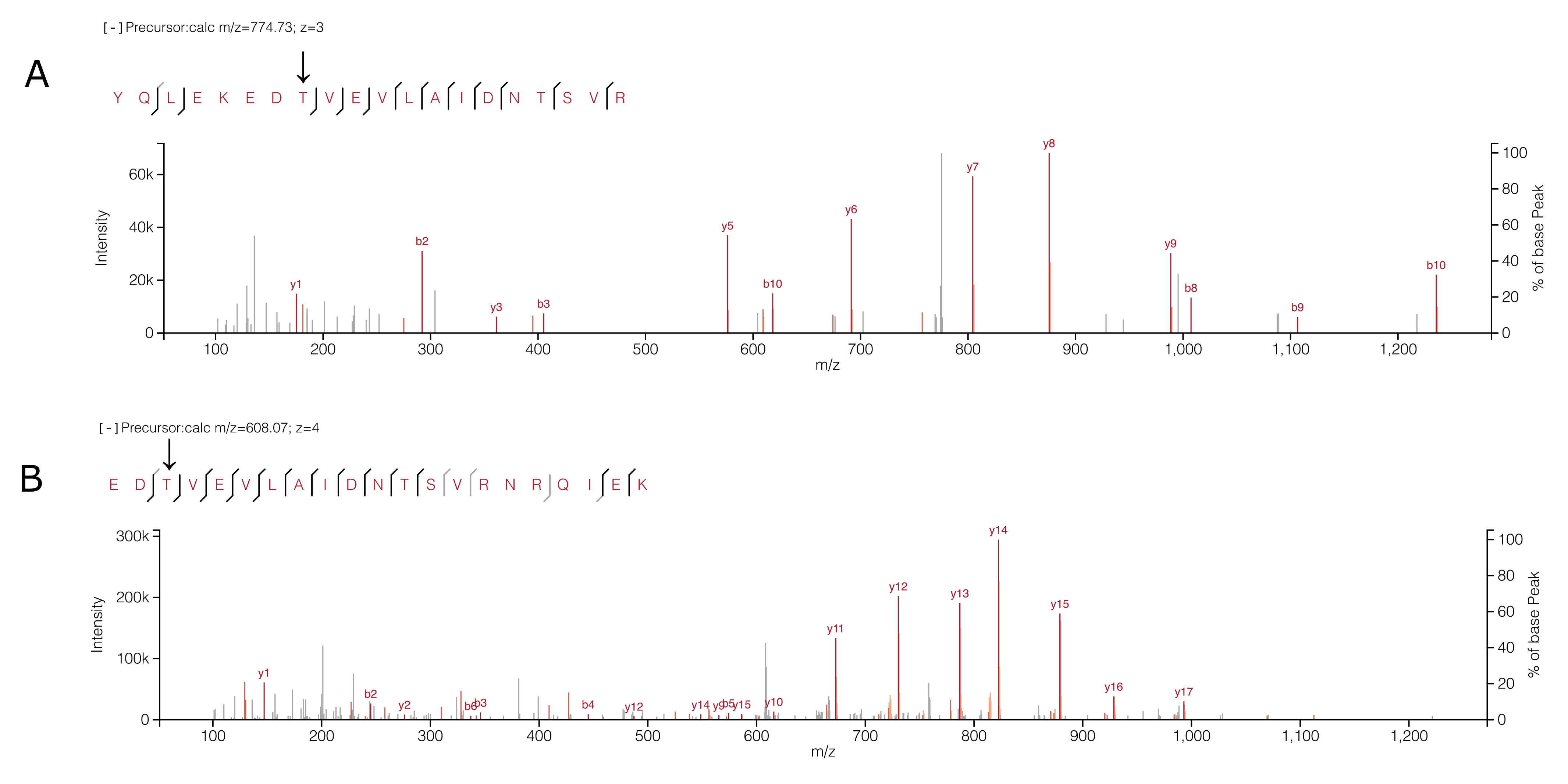

### Fig_S6.png

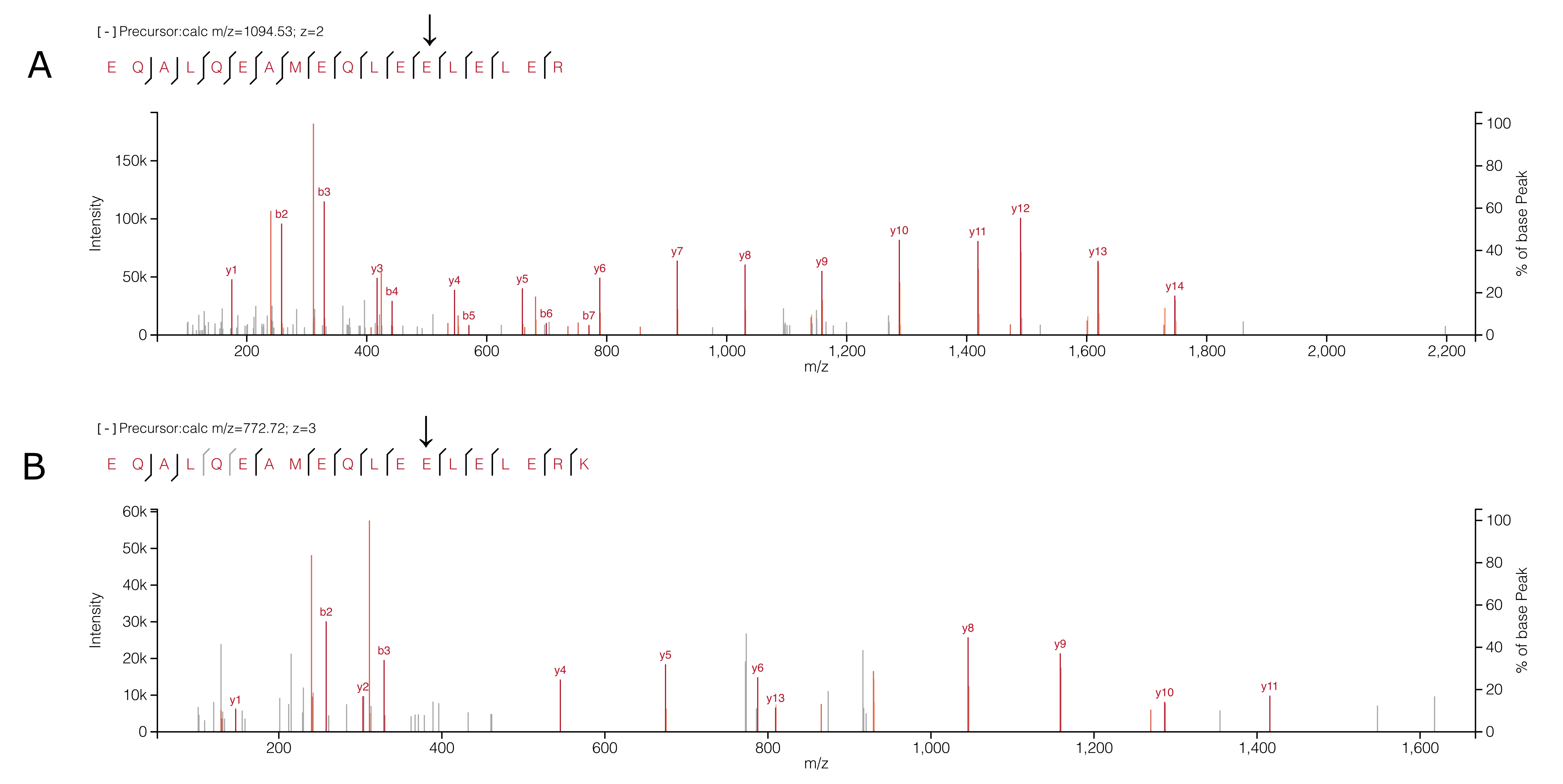
